# A root symbiont rewires tomato polyamine network to enhance herbivore resistance

**DOI:** 10.64898/2026.01.30.700540

**Authors:** Iván Fernández, Sara Comerón-Tabernero, Beatriz Romero-Rodríguez, Pedro López-Gómez, Ana Isabel González-Hernández, Alfonso Cornejo, María J. Pozo, Víctor Flors, Ainhoa Martínez-Medina

**Affiliations:** Laboratory of Molecular Agroecology, Estación Experimental del Zaidín - Consejo Superior de Investigaciones Científicas (EEZ-CSIC), Granada, Spain; Superior Polytechnic School of Zamora, University of Salamanca, Zamora, Spain; Institute for Advanced Materials and Mathematics, Department of Sciences, Public University of Navarre (UPNA), Pamplona, Spain; Department of Soil and Plant Microbiology, Estación Experimental del Zaidín - Consejo Superior de Investigaciones Científicas (EEZ-CSIC), Granada, Spain; Plant Immunity and Biochemistry Laboratory, Department of Biology, Biochemistry and Natural Sciences, Universitat Jaume I, Castellón de la Plana, Spain

**Keywords:** Endophyte, herbivore resistance, polyamine metabolism, root symbiosis, tomato defense, *Trichoderma* sp

## Abstract

Plants rely on specialized metabolites to defend against herbivores, yet how root-associated symbiotic microbes influence their regulation, coordination, and overall contribution to defense remains widely unknown. We hypothesized that symbiotic microbes can reprogram specialized metabolic networks, thereby enhancing herbivore resistance, with a particular focus here on polyamine metabolic network. Using the fungal symbiotic endophyte *Trichoderma harzianum*, tomato, and the herbivore *Spodoptera exigua*, we combined greenhouse bioassays with molecular, genetic, and metabolomic analyses to investigate how *Trichoderma* symbiosis reshapes the plant polyamine metabolic network, and affects plant-herbivore interactions. We found that in the absence of the root symbiosis, herbivory primarily activated polyamine uptake transport and catabolism, reflecting a stress-driven turnover response. *Trichoderma* symbiosis markedly reconfigured the polyamine network. Symbiosis primed uptake transport and catabolic responses, enhanced polyamine flux through activation of the ornithine decarboxylase pathway, and redirected polyamines into conjugated metabolites with anti-herbivore activity. Genetic analyses and dietary supplementation bioassays confirmed that this metabolic rewiring contributes to *Trichoderma*-induced resistance to herbivory, linking primary metabolic routes to the accumulation of specialized defense compounds. Our study highlights root symbionts as key modulators of plant metabolism, showing that specialized metabolite diversity can be shaped by symbiotic interactions.

## INTRODUCTION

Plants have coexisted and coevolved with insect herbivores for over 400 million years (Ehrlich & Raven, 1964; Labandeira & Currano, 2013). Over this vast evolutionary timespan, they have evolved a diverse arsenal of adaptive strategies to deter herbivory. Specialized metabolites including terpenoids, alkaloids, phenylpropanoids, fatty acid derivatives, and amino acid derivatives serve as pivotal chemical defenses in plant-herbivore interactions (Erb & Kliebenstein, 2020; Xu & Gaquerel, 2025). Their production relies on primary metabolic pathways that provide precursors, reducing power, and energy, resulting in tightly regulated biosynthesis (Li et al., 2024). Despite their recognized importance for plant fitness and survival, the ecological functions of many specialized metabolites in complex environments remain poorly understood (Huang & Dudareva, 2023), and the regulatory networks controlling their biosynthesis, transformation, and homeostasis are still largely unresolved (Kliebenstein, 2023).

Polyamines such as putrescine, spermidine, and spermine are involved in the regulation of critical plant processes, including growth, development and abiotic stress tolerance (Blázquez, 2024; Nawaz et al., 2026). More recently, they have further emerged as relevant regulators of plant defense against insect herbivory (Malik et al., 2014; Subramanyam et al., 2015; Zhang et al., 2023). These small nitrogen-containing molecules can act directly against herbivores and feed into the biosynthesis of diverse specialized defense compounds, linking primary and specialized metabolism (Kaur et al., 2010; Kerchev et al., 2012; Alamgir et al., 2016; Sivakumar et al., 2022; Zu et al., 2024). Polyamine levels are finely tuned through a dynamic network of biosynthesis, interconversion, transport, catabolism, and conjugation, enabling plants to adjust to developmental and environmental cues (Blázquez, 2024; Nawaz et al., 2026). In most plants, putrescine can be synthesized via the ornithine decarboxylase (ODC) or arginine decarboxylase (ADC) pathways providing metabolic flexibility (Cermanová et al., 2025). Downstream, spermidine and spermine are produced via aminopropyl transfer reactions, while catabolism mediated by diamine and polyamine oxidases (DAO and PAO, respectively) maintains homeostasis and generates signaling molecules such as hydrogen peroxide (H_2_O_2_) and γ-aminobutyric acid (GABA) (Salvi & Tavladoraki, 2020; Li et al., 2021). Conjugation with phenylpropanoids produces phenolamides, which contribute to anti-herbivore defense (Gaquerel et al., 2014). Although polyamines and their derivatives are increasingly recognized as hallmarks of plant defense, the roles of individual network components and the mechanisms regulating their activity under complex ecological contexts remain unclear.

Plants also share an ancient coevolutionary history with symbiotic microbes, including mycorrhizal fungi and plant growth-promoting rhizofungi and rhizobacteria (Delaux & Schornack, 2021). These interactions, widespread across terrestrial ecosystems, are increasingly recognized as major drivers of plant metabolic diversity (Delaux & Schornack, 2021; de Vries & Feussner, 2024). Root symbionts can prime local and systemic defenses, enhancing resistance to pathogens and herbivores through microbe-induced resistance (MIR, Pieterse et al., 2014). MIR often reshapes the plant metabolome, boosting specialized metabolites including phenolamides, alkaloids, phenylpropanoids, glucosinolates, and terpenoids, which contribute to herbivore toxicity (Pangesti et al., 2016; Papantoniou et al., 2021; Rivero et al., 2021, 2025; Abbas et al., 2024; Lv et al., 2024; Van Hee et al., 2025). However, despite evidence of MIR-driven metabolic reprogramming, how root symbiosis influences the evolution, regulation, and coordination of specialized metabolic networks remains unresolved.

Experimental evidence indicates that plant polyamine metabolism, including biosynthesis, accumulation and conjugation can be modulated by functionally diverse root-associated mutualists, such as mycorrhizal fungi, *Trichoderma* spp., and plant growth promoting rhizobacteria (Brotman et al., 2012; Salazar-Badillo et al., 2015; Coppola et al., 2019; Plett et al., 2020; Rivero et al., 2021; Nikhil et al., 2024; Minchev et al., 2026). However, it remains largely unexplored how microbial symbioses influence the overall functioning of the tightly regulated polyamine metabolic network, and how modulation of specific network components contributes to the plant’s adaptive responses to herbivory. We hypothesize that root symbiotic microbes adjust this coordinated network, enhancing the plant’s capacity to respond to herbivore attack. Using the MIR-inducing symbiotic fungus *Trichoderma harzianum* (hereafter Trichoderma), we aimed to understand whether and how root symbiosis reshapes the polyamine network and how its key functional elements contribute to enhanced resistance against herbivory. We employed tomato (*Solanum lycopersicum* L) and the herbivore *Spodoptera exigua* (Hübner, 1808), combining greenhouse bioassays with molecular, genetic, and metabolomic approaches.

Our study shows that Trichoderma root symbiosis primes herbivore-induced polyamine uptake transport and catabolism, while simultaneously activating de novo biosynthesis via the ODC pathway and driving conjugation into putrescine-derived phenolamides with anti-herbivore activity. These findings reveal that root symbionts not only amplify existing defense responses but can also restructure metabolic pathways, expanding the diversity and complexity of the specialized metabolite landscape under herbivore pressure. Our work provides a mechanistic framework for understanding how symbiotic microbes can fine-tune plant specialized metabolism to enhance adaptation and resistance in complex ecological contexts.

## RESULTS

### Trichoderma symbiosis reduces the performance of *Spodoptera exigua*

We first tested whether Trichoderma symbiosis modulates tomato resistance to *S. exigua*. In greenhouse bioassays, larvae fed on Trichoderma-inoculated plants gained significantly less weight compared to those on noninoculated plants (Figure 1A), whereas overall larval survival was not affected (Figure 1B). Notably, during the first 6 days of herbivory larvae showed a marginally lower probability of survival on Trichoderma-inoculated plants (Cox regression, days 1-3: exp(coef) = 4.271, *P*=0.08). Trichoderma inoculation did not affect plant growth (Supplementary Figure 1). These results indicate that Trichoderma symbiosis reduces the performance of *S. exigua*, primarily by impairing larval weight gain.

**Figure 1.**
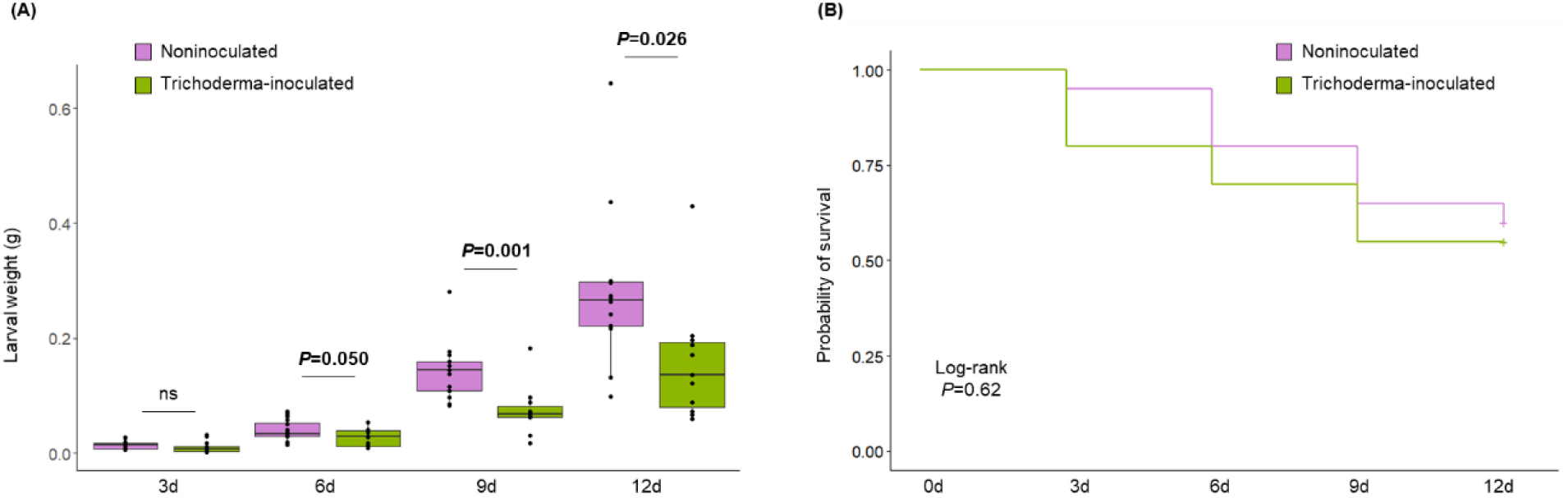
Impact of *Trichoderma harzianum* root colonization on *Spodoptera exigua* larval performance. (A) Larval weight and (B) survival probability were periodically assessed for *S. exigua* feeding on leaves from tomato plants root inoculated or not with *T. harzianum*. (A) Boxplots represent the interquartile range (IQR); horizontal lines indicate medians, whiskers indicate 1.5 times the IQR, and dots represent individual biological replicates (n = 15-20). *P* values (bold, *P* < 0.05) indicate results of *t*-tests between inoculated and noninoculated plants at each time point after infestation: 0d, 3d, 6d, 9d, and 12d. (B) Kaplan-Meier survival curves (n = 20 per treatment). Significant differences between treatments were determined using log-rank tests. The experiment was repeated three times, yielding similar results.

### Trichoderma symbiosis activates the ODC-branch of the polyamine biosynthesis pathway to induce herbivore resistance

Polyamine biosynthesis in tomato proceeds through two main branches: the ODC branch and ADC branch (Supplementary Figure 2). In the ODC branch, herbivory in noninoculated plants did not modify the transcript levels of *ORNITHINE DECARBOXYLASE 1/2* (*ODC1*, *ODC2*); and Trichoderma colonization alone, also had not effect on *ODC1*, *ODC2* expression (Figure 2A, Supplementary Table 1). By contrast, in Trichoderma-inoculated plants herbivory induced *ODC1* and *ODC2*, leading to significantly higher transcript levels compared to non-challenged plants (Figure 2A, Supplementary Table 1). Along similar lines, following herbivory, Trichoderma-inoculated plants displayed elevated ODC enzymatic activity (Figure 2B) and higher ornithine levels (Figure 2C), compared to noninoculated plants.

**Figure 2.**
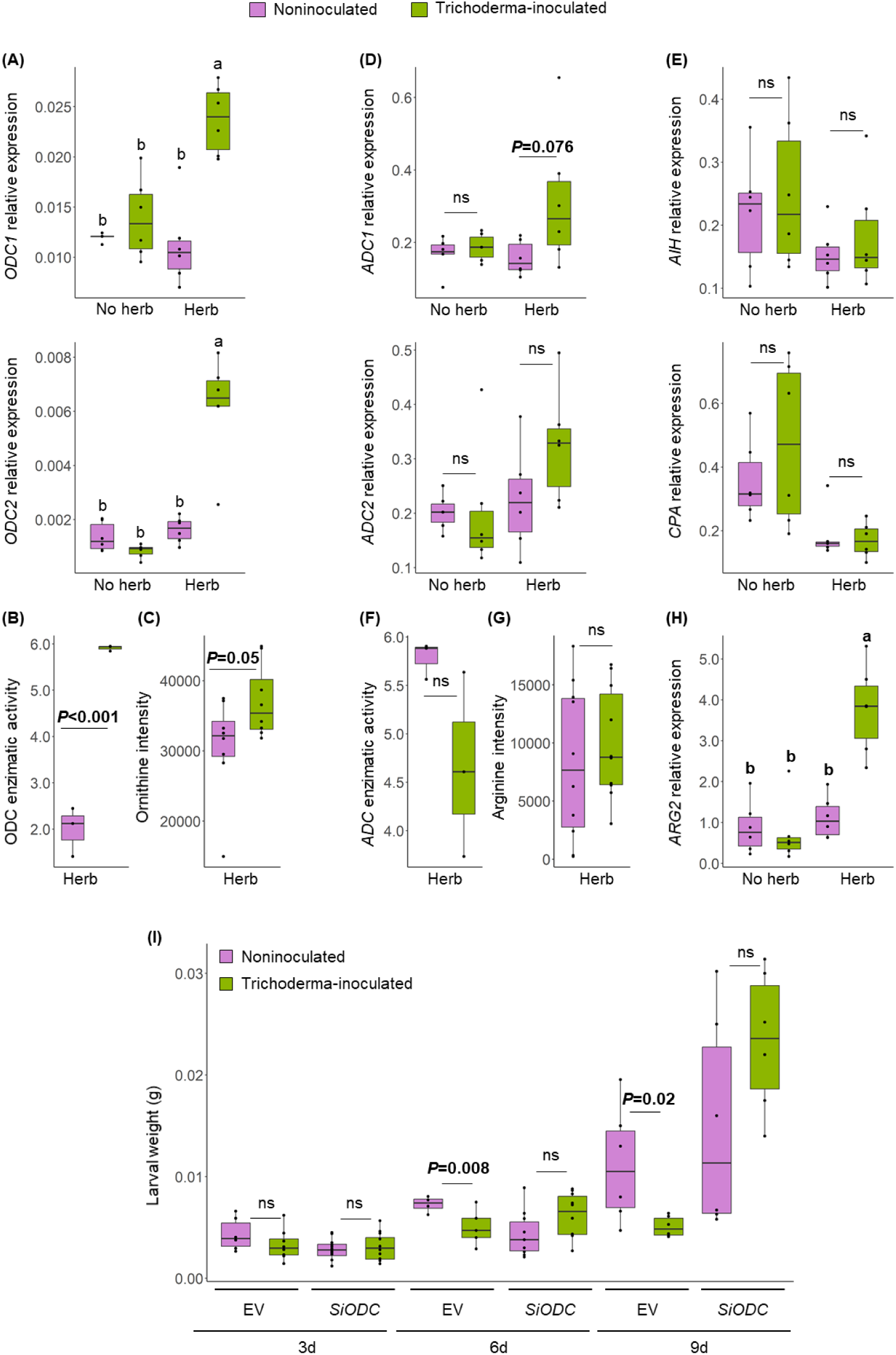
*Trichoderma harzianum*-mediated modulation of polyamine biosynthesis pathways. (A-C) Effect of *T. harzianum* root colonization on the ornithine decarboxylase (ODC) branch of polyamine biosynthesis: (A) transcript levels of *ORNITHINE DECARBOXYLASE 1* (*ODC1*) and *ORNITHINE DECARBOXYLASE 2* (*ODC2*); (B) ODC enzymatic activity; and (C) ornithine content. (D-H) Effect on the arginine decarboxylase (ADC) branch: (D) transcript levels of *ARGININE DECARBOXYLASE 1* (*ADC1*) and *ARGININE DECARBOXYLASE 2* (*ADC2*); (E) transcript levels of *AGMATINE IMINOHYDROLASE* (*AIH*) and *N-CARBAMOYLPUTRESCINE AMIDOHYDROLASE* (*CPA*); (F) ADC enzymatic activity; (G) arginine content; and (H) transcript levels of *ARGINASE 2* (*ARG2*). Plants were inoculated or not with *T. harzianum* and analyzed under control conditions or after 1 day of *Spodoptera exigua feeding*. (I) Role of the ODC branch in *Trichoderma*-induced resistance to herbivory. Larval weight of *S. exigua* was measured at 3, 6, and 9 days after infestation on tomato lines silenced for *ORNITHINE DECARBOXYLASE* (*SiODC*) and empty vector controls (EV), either inoculated or not with *T. harzianum*. Boxplots represent the interquartile range (IQR); horizontal lines indicate medians, whiskers extend to 1.5 × IQR, and dots represent individual biological replicates (n = 6-10). Different letters indicate significant differences (Tukey’s post hoc test, *P* < 0.05). *P* values indicate significant t-test comparisons (*P* < 0.05) between inoculated and non-inoculated plants for each genotype and/or condition. ns, non-significant. The experiments were repeated twice with similar results.

By contrast, the ADC branch was largely unaffected by Trichoderma. Neither herbivory nor Trichoderma inoculation significantly altered the expression of *ARGININE DECARBOXYLASE 1/2* (*ADC1*, *ADC2*) (Figure 2D, Supplementary Table 1). Similarly, transcripts of *AGMATINE IMINOHYDROLASE* (*AIH*) and (*N-CARBAMOYLPUTRESCINE AMIDOHYDROLASE* (*CPA*) (Figure 2E, Supplementary Table 1), ADC enzymatic activity (Figure 2F), and arginine content (Figure 2G) were unaffected. *ARG2* encodes a wound-inducible arginase that converts arginine into ornithine (Chen et al., 2024), supplying substrate for the ODC branch and potentially fueling ODC-mediated polyamine biosynthesis. We found that Trichoderma symbiosis enhanced *ARG2* expression upon herbivory (Figure 2H, Supplementary Table 1), whereas *ARG1* remained unchanged (Supplementary Figure 3). Together, these results indicate that in response to herbivory, Trichoderma symbiosis activates de novo polyamine biosynthesis, specifically through the ODC branch, a response not evident in noninoculated plants.

To assess the functional contribution of ODC-mediated polyamine biosynthesis to Trichoderma-induced resistance, we performed bioassays using a RNAi line impaired in the ODC branch (*SiODC*; González-Hernández et al., 2022). *T. harzianum* colonization was confirmed at comparable levels in the roots of silenced and EV plants (data not shown). In empty vector (EV) plants, Trichoderma significantly reduced larval weight gain, with effects evident at 6 and 9 days of herbivory (Figure 2I, Supplementary Table 1). This protective effect was completely abolished in the *SiODC* line, where larvae grew equally well on inoculated and noninoculated plants (Figure 2I, Supplementary Table 1). Altogether, these findings indicate that the ODC branch of the polyamine biosynthesis pathway is required for Trichoderma-induced resistance.

### Trichoderma symbiosis limits herbivory-induced free polyamine accumulation without altering polyamine interconversion

We next explored the impact of Trichoderma symbiosis and herbivory on free polyamine pool, including putrescine, spermidine and spermine. In noninoculated plants, herbivory did not affect putrescine but increased spermidine and spermine (Figure 3A, Supplementary Table 1). Strikingly, in Trichoderma plants, herbivory did not trigger free polyamine accumulation (Figure 3A, Supplementary Table 1). Moreover, following herbivory, a significantly lower accumulation of putrescine and spermidine was observed in Trichoderma plants compared to noninoculated plants (Figure 3A, Supplementary Table 1). Thus, the activation of polyamine biosynthesis by Trichoderma symbiosis was not accompanied by enhanced accumulation of free polyamines, suggesting downstream mobilization.

**Figure 3.**
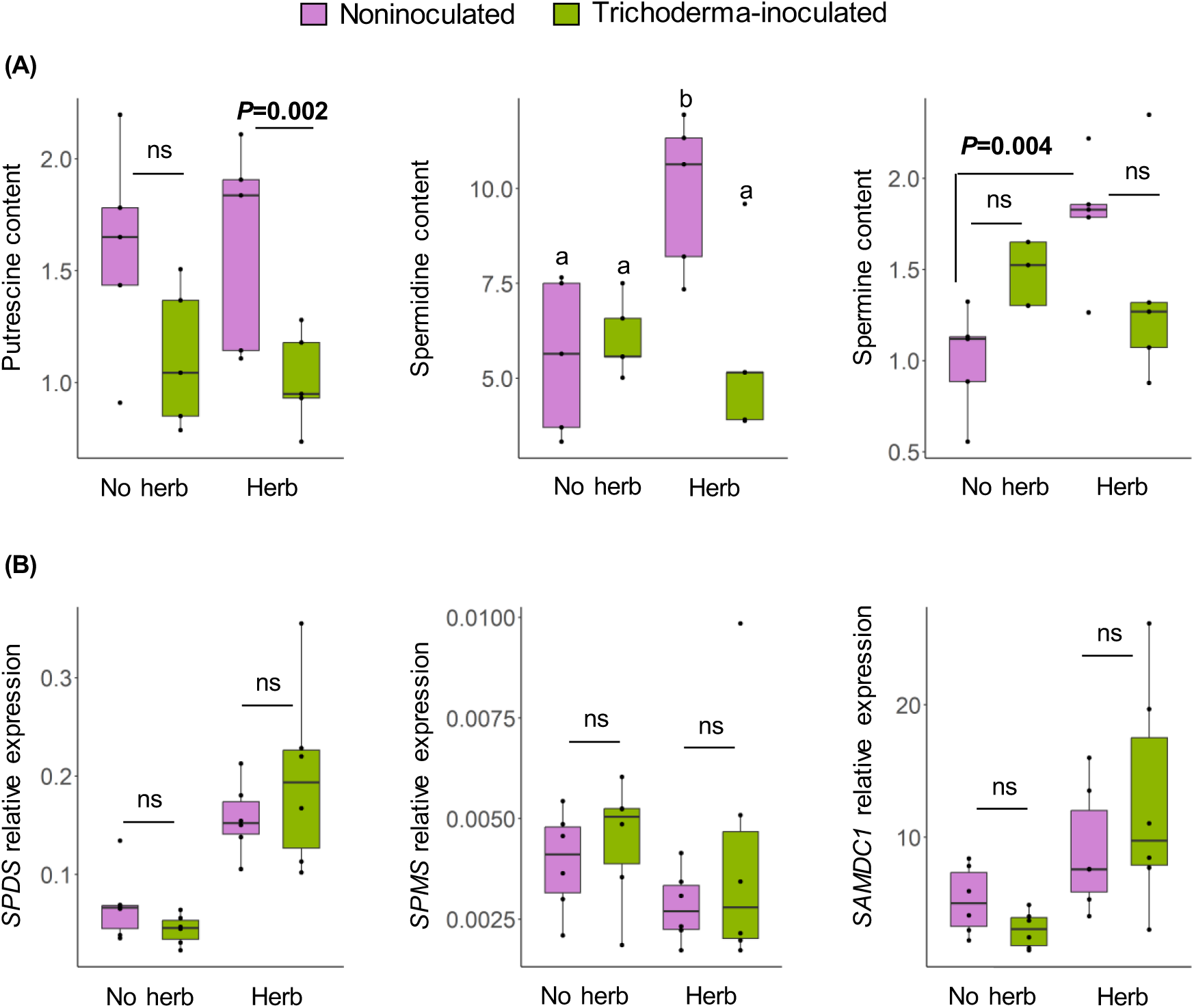
Impact of *Trichoderma harzianum* root colonization and *Spodoptera exigua* herbivory on free polyamine levels and interconversion. (A) Free polyamine content: putrescine, spermidine, and spermine. (B) Expression of polyamine interconversion-related genes: *SPERMIDINE SYNTHASE* (*SPDS*), *SPERMINE SYNTHASE* (*SPMS*), and *S-ADENOSYLMETHIONINE DECARBOXYLASE 1* (*SAMDC1*). Plants were inoculated or not with *T. harzianum* and analyzed under non-herbivory or 1-day herbivory conditions. Boxplots represent the IQR; horizontal lines indicate medians, whiskers 1.5 times the IQR, and dots individual biological replicates (n = 5-6). Different letters indicate significant differences (Tukey’s post hoc test, *P* < 0.05). *P* values indicate significant *t*-test results (*P* < 0.05). ns, non-significant. The experiment was repeated twice, yielding similar results.

Polyamine interconversion was not significantly altered by Trichoderma symbiosis. In noninoculated plants, herbivory upregulated *SPERMIDINE SYNTHASE* (*SPDS*, Factor Herbivory *P*<0.001, Supplementary Table 1), whereas *SPERMINE SYNTHASE* (*SPMS*) remained unaffected (Figure 3B, Supplementary Table 1). Herbivory also enhanced *S-ADENOSYLMETHIONINE DECARBOXYLASE 1* (*SAMDC1*, Factor Herbivory *P*<0.005, Supplementary Table 1) whose encoded enzyme provides aminopropyl donors for spermidine and spermine synthesis. Trichoderma inoculation did not significantly affect *SPDS*, *SPMS*, or *SAMDC1* (Figure 3B, Supplementary Table 1). These results indicate that during herbivory, Trichoderma symbiosis reduces the pool of free polyamines, specifically putrescine and spermidine, without affecting their interconversion, pointing to alternative metabolic fates.

### Trichoderma symbiosis enhances herbivore-induced polyamine transport and catabolism

We analyzed key genes related to polyamine transport and catabolism. In noninoculated plants, herbivory induced *PUTRESCINE UPTAKE TRANSPORTER 3* (*PUT3*). This effect was significantly amplified in Trichoderma-inoculated plants (Figure 4A, Supplementary Table 1), suggesting that Trichoderma symbiosis reinforces putrescine mobilization during herbivore attack. By contrast, expression of *DIAMINE OXIDASES 1/2/3* (*DAO1-3*) which are involved in putrescine catabolism producing H_2_O_2_ and GABA, was unaffected by either treatment (Figure 4B, Supplementary Table 1), indicating that this branch of putrescine turnover remained unchanged.

**Figure 4.**
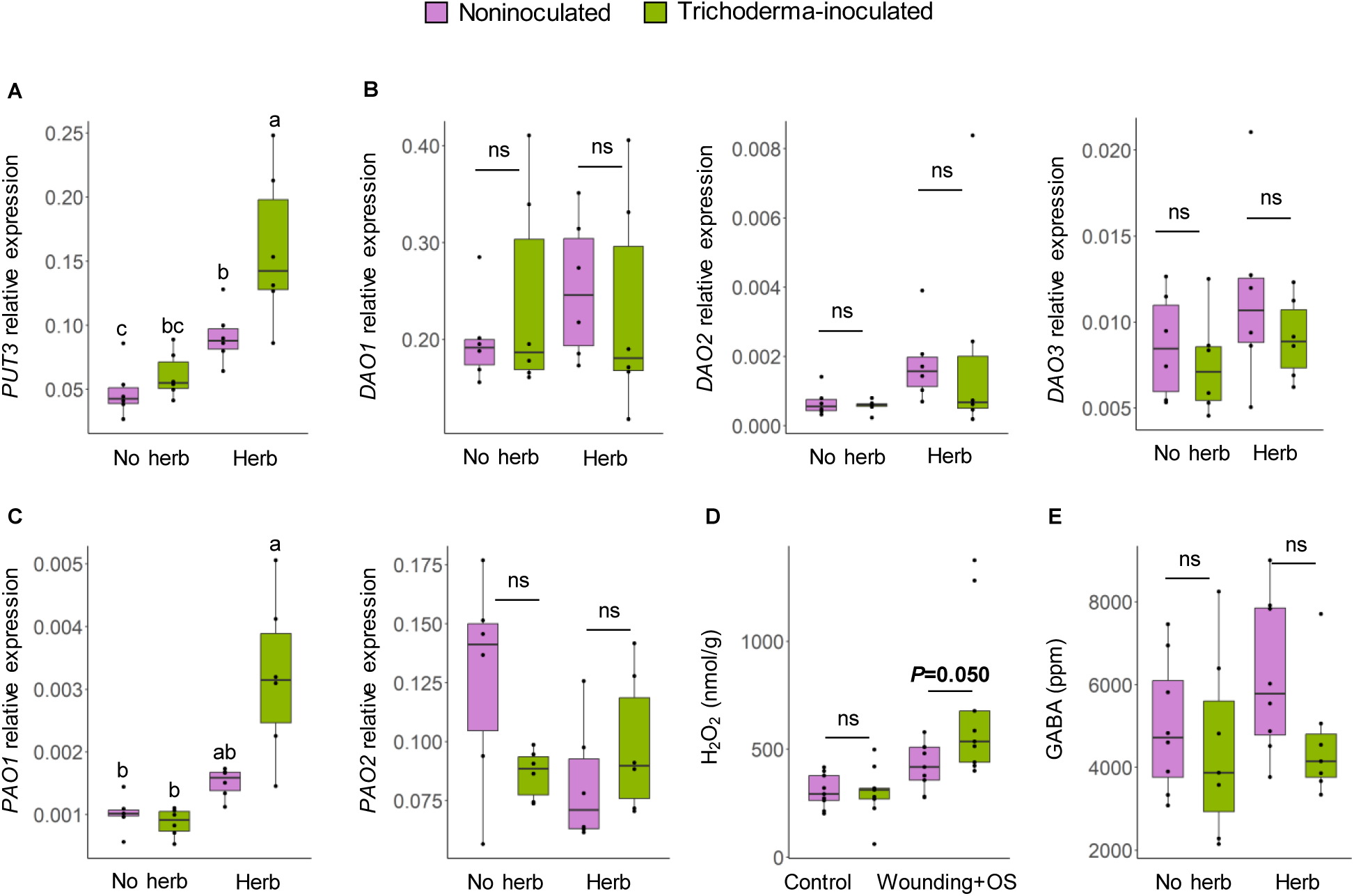
Effect of *Trichoderma harzianum* root colonization and *Spodoptera exigua* herbivory on polyamine transport and catabolism. Transcript levels of (A) *PUTRESCINE UPTAKE TRANSPORTER 3* (*PUT3*), (B) *DIAMINE OXIDASE 1/2/3* (*DAO1*, *DAO2* and *DAO3*), and (C) *POLYAMINE OXIDASES 1/2* (*PAO1* and *PAO2*). (D) Hydrogen peroxide levels, (E) amount of γ-aminobutyric acid (GABA). Plants were inoculated or not with *T. harzianum* and analyzed under non-herbivory conditions or after 1 day of *S. exigua* feeding (in A-C, E), or 20 min after simulated herbivory (wounding + oral secretion, OS, in D). Boxplots represent the IQR; horizontal lines indicate medians, whiskers 1.5 times the IQR, dots individual biological replicates (n = 6-10). Different letters indicate significant differences (Tukey’s post hoc test, *P* < 0.05). *P* values indicate significant *t*-test results (*P* < 0.05). ns, non-significant. The experiment was repeated twice, yielding similar results.

Herbivory increased the expression of *POLYAMINE OXIDASE 1* (*PAO1*, Factor Herbivory *P*<0.001, Supplementary Table 1) but not *POLYAMINE OXIDASE 2* (*PAO2*, Figure 4C, Supplementary Table 1). Notably, herbivore-triggered induction of *PAO1* was stronger in Trichoderma plants, compared to noninoculated plants, while Trichoderma did not affect *PAO2* (Figure 4C, Supplementary Table 1). In tomato, *PAO1* is triggered by wounding and functions as a spermidine and spermine oxidase, producing H₂O₂ (Hao et al., 2018). According with these expression patterns, an herbivory mimic treatment (mechanical wounding plus *S. exigua* oral secretions) triggered H₂O₂ production (Factor Herbivory *P*< 0.001) which was significantly higher in Trichoderma-inoculated plants (Figure 4D, Supplementary Table 1). Consistent with the lack of DAO activation, GABA levels were not affected by either herbivory or Trichoderma treatment (Figure 4E, Supplementary Table 1), further reinforcing that DAO-mediated putrescine catabolism was not activated by either herbivory or Trichoderma symbiosis. Therefore, our results indicate that Trichoderma symbiosis enhances herbivore-induced *PUT3* expression and *PAO1*-mediated catabolism, likely redirecting free polyamines, particularly spermidine and/or spermine, toward oxidative turnover products, including H₂O₂.

### Trichoderma symbiosis promotes the accumulation of putrescine-derived phenolamides with antiherbivore activity

Polyamine conjugation with hydroxycinnamic acids represents a key metabolic branch regulating polyamine homeostasis. In noninoculated plants, herbivory specifically induced the expression of *PUTRESCINE HYDROXYCINNAMOYL TRANSFERASE 2* (*PHT2*) but not *PUTRESCINE HYDROXYCINNAMOYL TRANSFERASE 1/3* (*PHT1*, *PHT3*), or SPERMIDINE HYDROXYCINNAMOYL TRANSFERASE (*SHT*) (Figure 5A, Supplementary Figure 4, Supplementary Table 1). In tomato, an additional member of this family, *PUTRESCINE HYDROXYCINNAMOYL TRANSFERASE 4* (*PHT4*), is present, but it appears transcriptionally inactive in foliar tissue under the conditions tested. Strikingly, in Trichoderma-inoculated plants, herbivory elicited a stronger induction of both *PHT1* and *PHT2* compared to noninoculated plants (Figure 5A). Moreover, Trichoderma symbiosis enhanced *PHT1* expression even in the absence of herbivory, whereas *PHT3* and *SHT* remained unaffected (Figure 5A, Supplementary Figure 4, Supplementary Table 1).

**Figure 5.**
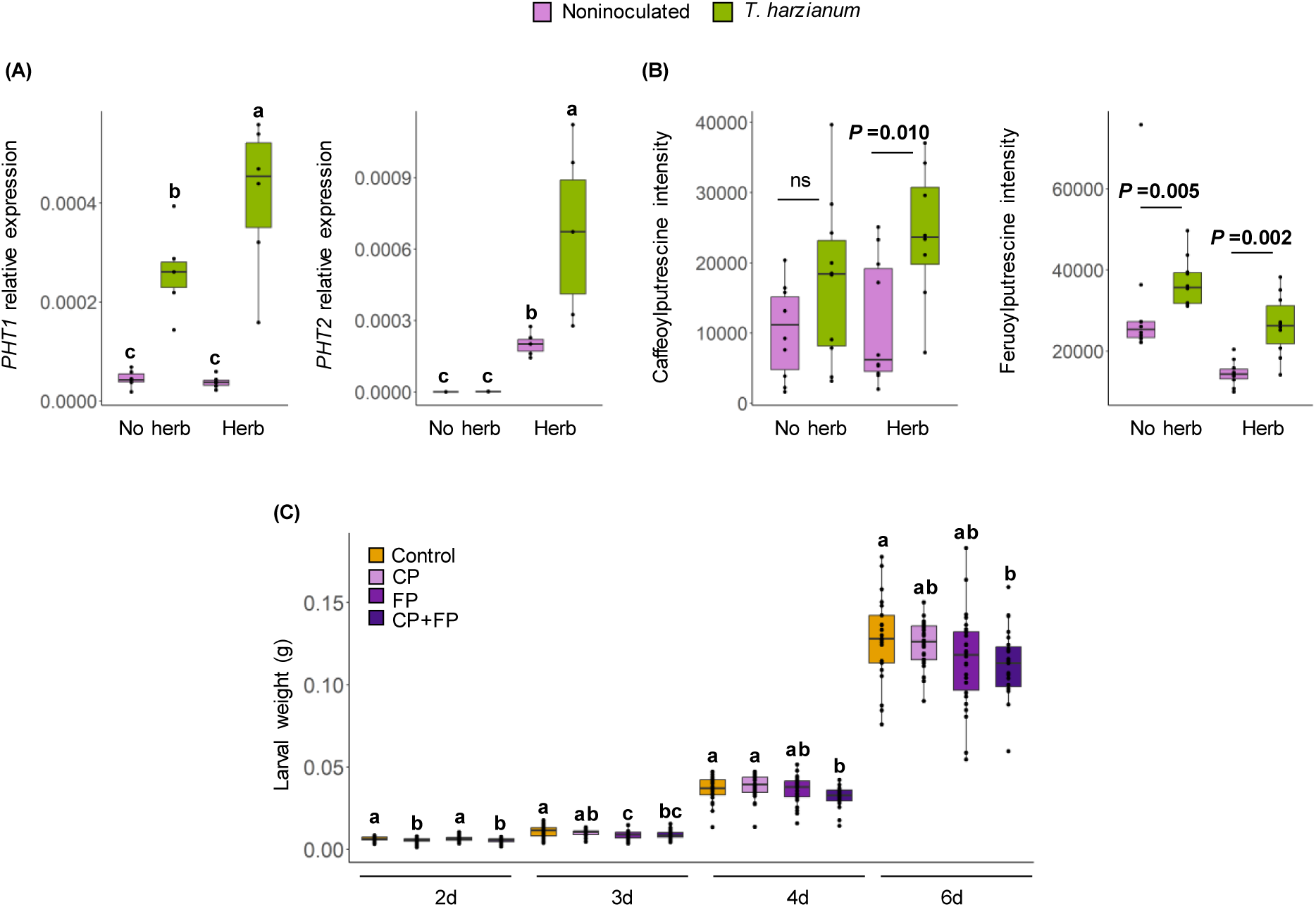
Impact of *Trichoderma harzianum* root colonization and *Spodoptera exigua* herbivory on the biosynthesis of antiherbivore phenolamides. (A) Transcript levels of polyamine conjugation-related genes *PUTRESCINE HYDROXYCINNAMOYLTRANSFERASE 1/2* (*PH1*, *PHT2*). (B) Intensity level of the polyamine conjugates caffeoylputrescine and feruloylputrescine from leaf samples analyzed by liquid chromatography-mass spectrometry (LC-MS). Plants were inoculated or not with *T. harzianum* and analyzed under non-herbivory conditions or after 1 day (A) and 2 days (B) of *S. exigua* feeding. Boxplots represent the interquartile range (IQR); horizontal lines indicate medians, whiskers 1.5 times the IQR, and dots individual biological replicates (n = 6-10). Different letters indicate significant differences according to Tukey’s post hoc test (*P* < 0.05). *P* values (bold, *P* < 0.05) indicate *t*-test results between inoculated and control plants. ns, non-significant. (C) Direct effects of caffeoylputrescine (CP) and feruloylputrescine (FP) in *S. exigua* larval growth. Larvae were fed on artificial diets supplemented with CP, FP, or a mixture of both (CP+FP). Control larvae were fed acidified Milli-Q water, corresponding to the solvent used. Box plots represent the interquartile range (IQR), the bisecting line represents the median, the whiskers represent 1.5 times the IQR, and the dots represent data points (n= 25 - 30 biological replicates). Different letters at each time point indicate statistically significant differences among treatments according to Tukey’s post hoc test (*P* < 0.05).

Metabolite profiling identified two main polyamine conjugates: caffeoylputrescine (CP) and feruloylputrescine (FP), that were confirmed by standards (Figure 5B, Supplementary Figure 4). We further identified feruloyldicumaroylspermidine, putatively annotated based on MS fragmentation patterns (Supplementary Figure 4). In noninoculated plants, herbivory did not significantly alter CP levels, but decreased FP accumulation (Factor Herbivory *P* < 0.001; Figure 5B; Supplementary Table 1). Remarkably, upon herbivore challenge, Trichoderma symbiosis led to enhanced accumulation of CP and FP compared with noninoculated plants. Furthermore, Trichoderma increased FP levels even in the absence of herbivory (Figure 5B).

To explore the direct impact of CP and FP on *S. exigua* performance, we conducted a bioassay using artificial diet supplemented with the pure compounds. Application of either CP or FP alone resulted in a transient reduction in larval weight (Figure 5C, Supplementary Table 1). Specifically, CP supplementation reduced larval weight after 2 days of feeding, whereas this effect was no longer evident at later time points (days 4 to 6; Figure 5C). In the case of FP, larval weight reduction was observed at 3 days, with no significant effects detected at the remaining days of the assay. Remarkably, the combined application of CP and FP led to a sustained and significant reduction in larval weight throughout the entire bioassay (Figure 5C, Supplementary Table 1). Together, these results demonstrate that Trichoderma symbiosis activates the conjugation branch of the polyamine metabolic network, promoting both constitutive and herbivore-induced accumulation of putrescine-derived phenolamides, particularly CP and FP, with antiherbivore activity.

### Tomato biosynthesis of putrescine-derived phenolamides contributes to herbivore resistance

We next aimed to assess whether PHT1/2 are indeed involved in the biosynthesis of putrescine-derived phenolamides and in tomato resistance to *S. exigua*. With this aim, we used virus-induced gene silencing (VIGS) to suppress *PHT1/2* expression using two different constructs, targeting two different *PHT1/2* regions (i.e.: target 1 and target 2). Notice that, due to the high sequence similarity between *PHT1* and *PHT2*, both genes were silenced simultaneously (Figure 6 A,D). We found that TRV::PHT1/2 plants, including either target 1 and target 2, accumulated significantly less FP after herbivory compared with TRV::00 controls, while CP was undetectable in these samples (Figure 6 B,E). Larvae feeding on TRV::PHT1/2 plants, using either of the two constructs targeting different regions, showed significantly greater weight gain at 6 days after feeding, and a marginally (0.05 < *P* < 0.1) higher gain at 9 days (Figure 6 C,F). Together, these findings demonstrate that PHT1/2-mediated biosynthesis of putrescine-derived phenolamides, specially FP, contributes to herbivore resistance. Moreover, they support the view that Trichoderma-induced activation of this conjugation pathway constitutes a mechanistic component of MIR.

**Figure 6.**
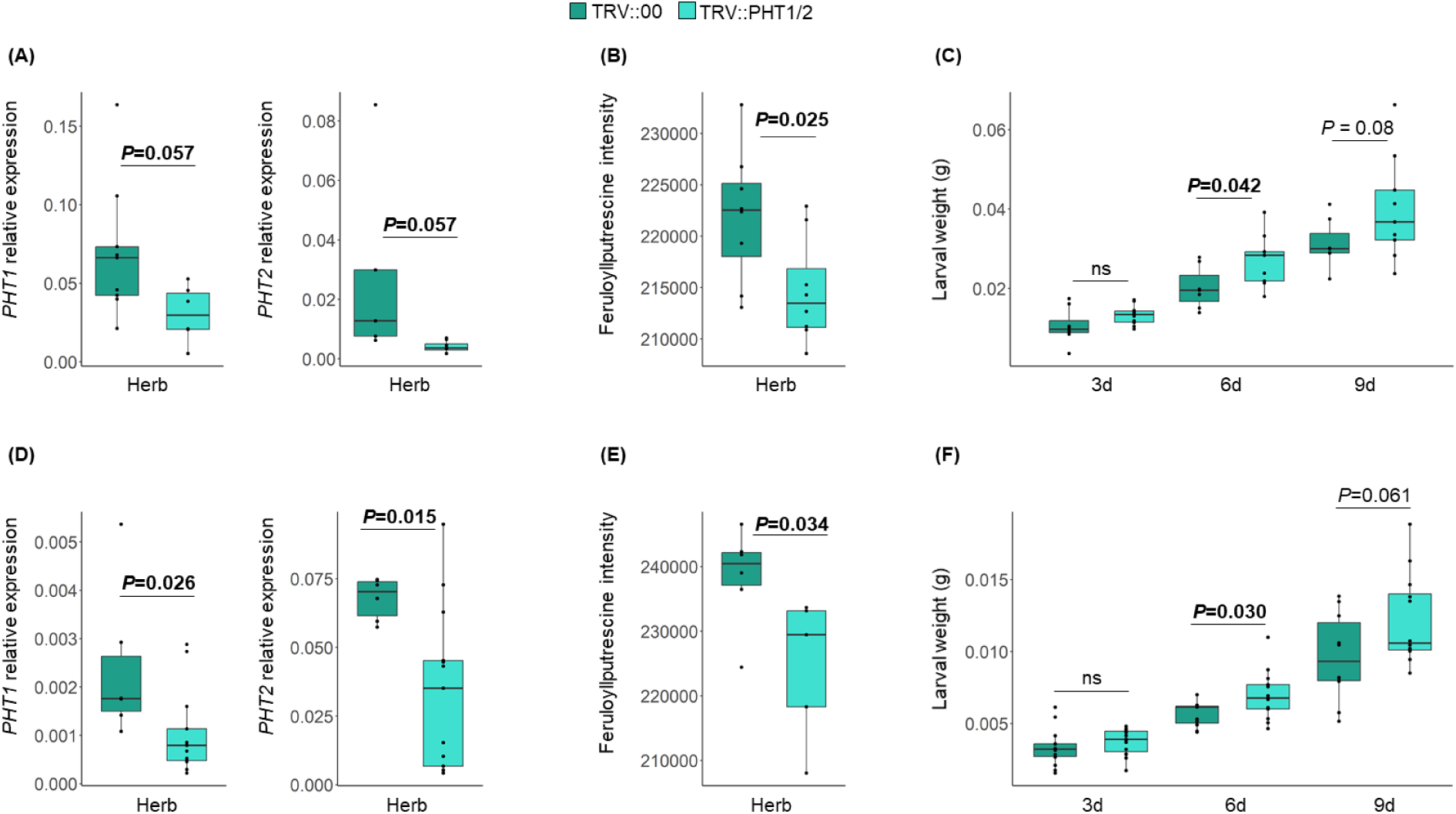
Role of putrescine-derived phenolamides biosynthesis in plant resistance to herbivory. Virus-induced gene silencing (VIGS) was used to suppress *PUTRESCINE HYDROXYCINNAMOYLTRANSFERASE 1*/2 (*PHT1/2*) using two different constructs targeting two different regions of *PHT1/2* (target 1, top panels; target 2, bottom panels). (A, D) Transcript levels of *PHT1* and *PHT1* in TRV control plants (TRV::00) and plants silenced with TRV::PHT1/2 constructs targeting region 1 (A) or region 2 (D). (B, E) Leaf feruloylputrescine (FP) accumulation in TRV controls and *PHT1/2*-silenced plants (region 1, B; region 2, E). (C, F) Weight of *Spodoptera exigua* larvae feeding on TRV controls and *PHT1/2*-silenced plants (region 1, C; region 2, F). Boxplots represent the interquartile range (IQR); horizontal lines indicate medians, whiskers 1.5 times the IQR, and dots correspond to individual biological replicates (n = 8-15). *P* values (bold, *P* < 0.05) indicate *t*-test results between treatments. ns, non-significant.

## DISCUSSION

Root symbionts are increasingly recognized as modulators of polyamine metabolism in their host plants, influencing biosynthesis, accumulation, and conjugation (Brotman et al., 2012; Salazar-Badillo et al., 2015; Coppola et al., 2019; Plett et al., 2020; Rivero et al., 2021; Nikhil et al., 2024; Minchev et al., 2026). However, despite these advances, how symbiotic associations rewire the integrated functioning of the polyamine metabolic network, and how such network-level adjustments shape plant defensive phenotypes, remains unresolved. Polyamine metabolism operates as a highly dynamic and interconnected system, coupling biosynthesis, transport, catabolism, and conjugation to generate a wide array of specialized metabolites. Through this organization, the polyamine network functions as a central regulatory hub linking primary metabolism with chemical defense and broader stress-adaptation processes (Blazquez, 2024). Consistent with our hypothesis, our findings show that the root symbiont *Trichoderma harzianum* reconfigures the tomato polyamine metabolic network, at multiple levels, enhancing resistance to herbivory through coordinated regulation of its major functional branches. Trichoderma symbiosis boosts herbivore-induced uptake transport-catabolism module, activates de novo polyamine biosynthesis via ODC-dependent pathway, and stimulates conjugation with hydroxycinnamic acids to produce phenolamides with anti-herbivore activity. Together, these responses show that the symbiotic fungus not only primes herbivore-triggered polyamine responses, but also engages additional metabolic branches, thereby increasing the complexity and diversity of the polyamine network, expanding thus the metabolic landscape of the host under herbivore pressure.

In the absence of the root symbiont, herbivory primarily activated the polyamine network branch associated with uptake transport (*PUT3*), spermidine synthesis (*SPDS*), and catabolism (*PAO*), leading to H₂O₂ accumulation and reflecting a general stress-induced turnover response (Figure 7). Polyamine oxidation and the resulting reactive oxygen species have been implicated in plant resistance to a variety of stresses, including herbivory (Yoda et al., 2006; Yang et al., 2024; Zu et al., 2024). In non-colonized plants, these polyamine-mediated responses likely reflect a general oxidative stress signaling response rather than a defense program specifically directed against herbivores. Remarkably, Trichoderma symbiosis strongly amplified this herbivore-induced module, enhancing *PUT3* expression and promoting *PAO1* induction, accompanied by increased H₂O₂ accumulation (Figure 7). By contrasts, *DAO* genes remained unchanged and GABA levels were unaffected by either herbivory or Trichoderma symbiosis, indicating that DAO-mediated putrescine catabolism was not activated under these conditions. These findings indicate that Trichoderma specifically primes PAO-driven oxidative polyamine turnover, likely redirecting free spermidine and/or spermine toward H₂O₂ production, consistent with the observed reduction in spermidine levels, and contributing to herbivory-induced oxidative signaling. Indeed, Trichoderma-mediated resistance has been linked to enhanced redox signaling and controlled production of reactive oxygen species (ROS), which function both as antimicrobial and anti-herbivore agents and as central signals in defense regulation (Contreras-Cornejo et al., 2011; Kerchev et al., 2012; Kumar et al., 2023; Mahanta et al., 2025). While symbiont-triggered ROS can originate from multiple enzymatic sources, our results indicate that the primed polyamine uptake-catabolism module via PAO, contribute to H₂O₂ dynamics during herbivory, although dedicated functional validation is still required. Notably, recent work shows that PUT3 and several polyamine-modifying enzymes form a coordinated regulatory module within the same genomic cluster in tomato, supporting the emerging view that polyamine turnover and H₂O₂ homeostasis constitute an integrated stress-response node (Yang et al., 2024).

**Figure 7.**
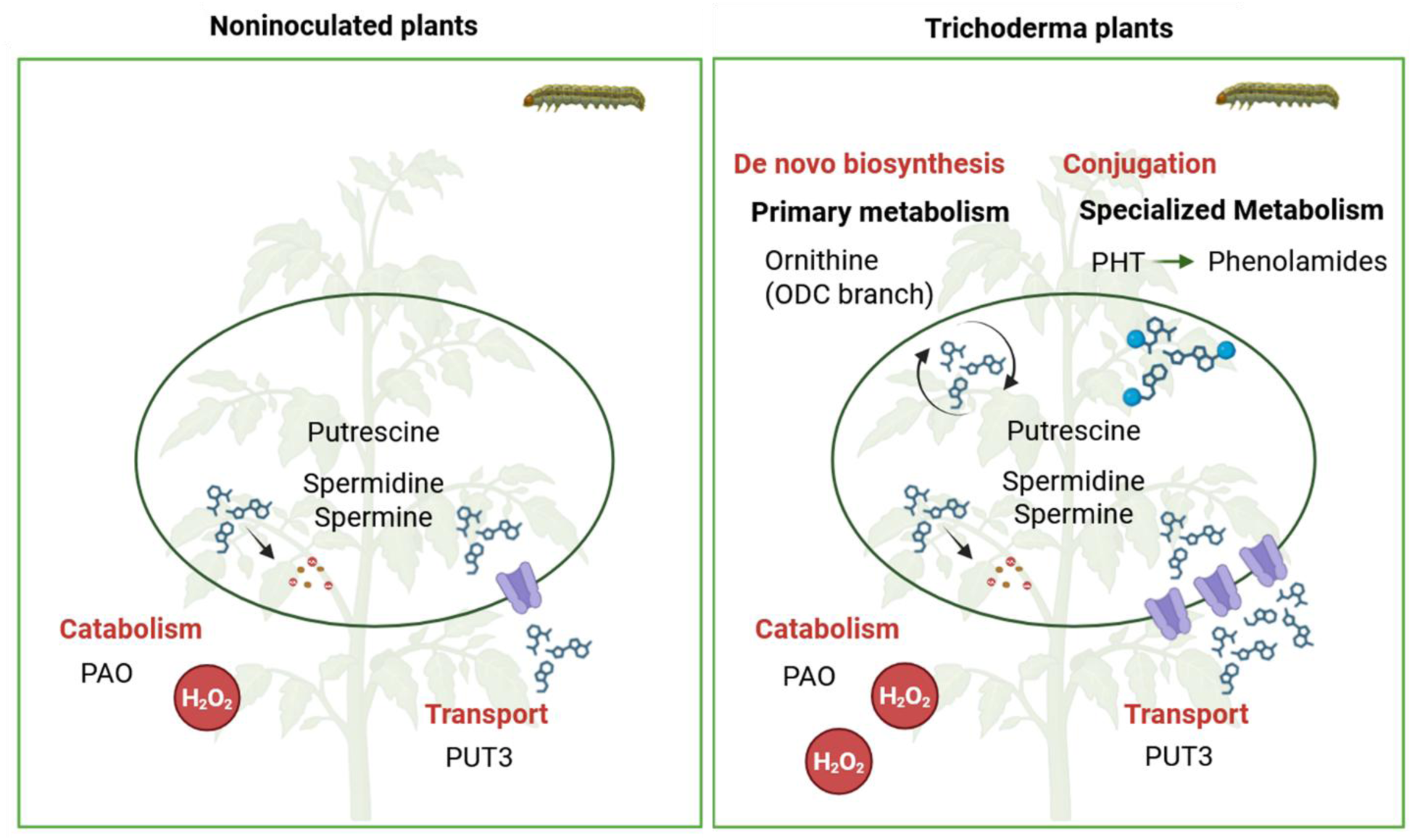
Trichoderma root symbiosis reconfigures the tomato polyamine network to enhance herbivore resistance. In plants lacking endophytic symbiosis, herbivory by *Spodoptera exigua* primarily activates the polyamine network branch associated with transport uptake (via PUT3) and catabolism (via PAO), resulting in hydrogen peroxide (H₂O₂) accumulation. This oxidative burst functions both as a direct toxic defense against herbivores and as a signaling cue for defense activation. (B) In *Trichoderma*-colonized plants, symbiosis primes polyamine transport and catabolic nodes, resulting in a controlled increase in H₂O₂ production that contributes to enhanced resistance. Following herbivory, symbiosis further activates *de novo* polyamine biosynthesis through the ODC-mediated pathway and redirects free polyamines, via PHT activation, toward the production of putrescine-derived phenolamides with antiherbivore activity. Overall, Trichoderma symbiosis rewires the polyamine metabolic network, functionally linking primary metabolism to specialized defense pathways, thereby increasing the complexity, diversity, and functional output of the polyamine-derived metabolome.

Even more strikingly, Trichoderma symbiosis reconfigured the polyamine network beyond the uptake transport and catabolism modules, activating both the biosynthetic and conjugation branches, which seemed to remain largely quiescent upon herbivory in plants lacking the symbiosis (Figure 7). This qualitative reprogramming guided our subsequent genetic analyses toward these specific pathway components. Upon herbivory, Trichoderma specifically induced the ODC pathway, as evidenced by elevated *ODC* transcript levels, increased ornithine accumulation, and higher ODC enzymatic activity. Notably, ornithine levels rose despite enhanced ODC activity, indicating that the increase was fueled by greater precursor availability rather than reduced consumption. Consistent with this, the symbiont upregulated *ARG2*, a jasmonic acid- and wound-responsive arginase that converts arginine into ornithine, thereby sustaining the ODC-driven biosynthetic flux (Chen et al., 2004). Functional assays confirmed the central role of this pathway in symbiont-mediated resistance: silencing *ODC* abolished Trichoderma-induced protection. Although direct metabolic flux measurements were not performed, our data indicate that Trichoderma symbiosis recruits the ODC pathway as a metabolic switch, channeling primary amino acid precursors into defense-oriented polyamine biosynthesis during herbivore attack.

Concomitant with the activation of the biosynthetic module, Trichoderma symbiosis also elicited the conjugation module of the polyamine network, enhancing the formation of putrescine-derived phenolamides (Figure 7). These specialized metabolites are generated through the conjugation of hydroxycinnamic acids with polyamines via acyltransferases, a process described in several plant species (Luo et al., 2009; Roumani et al., 2021; Bernard et al., 2022). In tomato, four putative hydroxycinnamoyltransferases (PHTs) have recently been identified (Roumani et al., 2025), yet their role in antiherbivore defense remained unexplored, and our understanding of phenolamide metabolism and regulation is still incomplete. Here, we found that Trichoderma enhanced *PHT1* and *PHT2* expression and promoted the accumulation of CP and FP. Bioassays with pure compounds revealed that both phenolamides were associated with reduced larval growth, with the strongest and most consistent effects observed when CP and FP were applied together. Consistent with this, putrescine-derived phenolamides have previously been linked to antiherbivore activity in other systems (Kaur et al., 2010; Minchev et al., 2026). Genetic manipulation further supported the defensive role of this conjugation pathway: simultaneous silencing of *PHT1* and *PHT2* in tomato reduced FP levels and promoted larval growth. Although the inducibility and defensive role of phenolamides vary across plant-insect systems (Kim et al., 2011; Gaquerel et al., 2014; Li et al., 2020; Kundu, 2023), our findings demonstrate that symbiosis-driven activation of the conjugation branch of the polyamine network constitutes a functionally relevant mechanism contributing to enhanced resistance against herbivory.

Noticeably, in non-colonized plants, herbivory did not trigger phenolamide accumulation, suggesting that activation of this branch is symbiosis-dependent. Phenolamide accumulation can be triggered by herbivory in several plants species (Pearce et al., 1998; Kaur et al., 2010; Alamgir et al., 2016). Yet, in certain plant-insect systems, this response is weak or absent and does not necessarily confer greater resistance emphasizing its context-dependent nature (Roumani et al., 2021). Consistent with our observations, Minchev et al. (2026) found that *Tuta absoluta* induced FP accumulation only in mycorrhizal plants, but not in plants lacking the symbiosis. The reasons and mechanistic basis for the lack of phenolamide induction by *S. exigua* in tomato are currently unknown, but may involve *Spodoptera*-mediated suppression of host immunity (Chen et al., 2019) and/or nitrogen allocation trade-offs that occurs during herbivory (Matsuno et al., 2009; Ullmann-Zeunert et al., 2013). Polyamine biosynthesis is a metabolically demanding process, particularly under stress conditions, and competes with other metabolic pathways for essential resources, including amino acids (Nawaz et al., 2026). In this context, Trichoderma-induced increase in ornithine may support both enhanced ODC-mediated polyamine biosynthesis and phenolamide formation, alleviating nitrogen constraints that might otherwise limit these defenses.

Overall, our study demonstrates that a root symbiont orchestrates a coordinated reprogramming of the tomato polyamine network, linking primary metabolism to specialized metabolite production and antiherbivore defense (Figure 7). The symbiont not only primes the uptake transport-catabolism module but also activates ODC-driven biosynthesis and phenolamide conjugation, thereby increasing the complexity, diversity, and functional contribution of the polyamine metabolomic network. Remarkably, these responses reveal that under stress, microbial symbionts can induce metabolic branches that seems to remain quiescent in the absence of the symbiont. Exploring this idea is particularly important because it challenges the prevailing view that diversification of specialized metabolites occurs predominantly as a direct response to biotic threats (Speed et al., 2015). By showing that symbiotic interactions can expand and reshape metabolic networks, our findings provide a mechanistic framework for understanding how root symbiosis dynamically fine-tune specialized metabolism to enhance plant adaptation and resistance under herbivore pressure.

## MATERIAL AND METHODS

### Plant, fungal and insect material

We used tomato (*Solanum lycopersicum*) cultivar Moneymaker. We further used the RNA interference (RNAi) transgenic line *SiODC* (González-Hernández et al., 2022), and the corresponding parental Moneymaker line carrying the empty vector pBIN19-RNAi. Seeds were kindly provided by the Biochemistry and Biotechnology Group (Universitat Jaume I, Castellón, Spain). The root endophytic fungus *Trichoderma harzianum* was maintained on potato dextrose agar at 4°C as part of our laboratory collection. The *T. harzianum* inoculum was prepared according to Martinez-Medina et al. (2009). Batches from the same *Spodoptera exigua* (Lepidoptera: Noctuidae) laboratory colony were used for the different experiments (kindly provided by Prof. Herrero, Valencia University, Spain). Larvae were reared with artificial diet (Hoffman et al., 1966) in a growth chamber maintained at 23 °C, 12-h light and 12-h dark cycle, and 50 % relative humidity.

### Plant growth and fungal inoculation and quantification

Tomato seeds were surface-sterilized in 5% NaClO4, rinsed thoroughly with sterile water, and germinated for 10 days in sterilized vermiculite at 25°C. Tomato seedlings were then transferred to 600 mL pots containing a sterile 1:1 mixture (v : v) of sand and vermiculite. They were assigned to one of the treatment groups: noninoculated plants, or plants inoculated with *T. harzianum*. Inoculation with *T. harzianum* was achieved by mixing the inoculum with the substrate to a final density of 1 × 10^6^ conidia g^-1^ before transplanting (Papantoniou et al., 2021). The plants were then placed in a completely randomized design in a glasshouse compartment with a 24 ± 3°C, 16-h light and 8-h dark cycle, and 60% relative humidity. The plants were watered three times a week: 2 days using tap water and 1 day using Hoagland nutrient solution (Hoagland & Arnon, 1950). Five weeks after transplanting, the plants were used for experiments. Before using the plants, we confirmed that *T. harzianum* had efficiently colonized the potting substrate of the corresponding treatments by using the plate count technique and potato dextrose agar amended with 50 mg L^-1^ rose bengal, and 100 mg L^−1^ streptomycin sulphate according to Papantoniou et al. (2021). The number of *T. harzianum* CFU in the potting media of *T. harzianum*-inoculated plants was similar to the initial inoculation values (i.e., 1 × 10^6^ conidia g^-1^). *T. harzianum* endophytic colonization was confirmed in surface-sterilized roots for all the tomato lines according to Martínez-Medina et al. (2017), and using the specific primers Tef1-F: 5′-GGTACTGGTGAGTTCGAGGCTG-3′ and Tef1-R: 5′- GGGCTCGATGGAGTCGATAG3′, which are specific for the constitutively expressed Trichoderma translation elongation factor-1α gene.

### Virus-induced *PHT1/PHT2* silencing

Tobacco rattle virus (TRV)-based silencing vectors pTRV1 and pTRV2::00 (empty vector) were kindly provided by Dr. Rodríguez Bejarano (University of Málaga, Spain). To simultaneously silence *PHT1* and *PHT2* genes, a 302 bp gene-specific fragment was amplified using a GeneScript plasmid containing the full-length *PHT2* sequence (pUC57_SlPHT2) as a template (GenBank: ON248951.1; Solgenomics: Solyc11g071480). Because of the high sequence similarity between *PHT1* and *PHT2* genes, specific silencing of a single gene was not feasible, resulting in simultaneous silencing of both genes. Target selection was performed using the VIGS tool available on the Sol Genomics Network (http://vigs.solgenomics.net) choosing two different regions (target 1 and target 2) with high distributions of n-mers in both genes and none off-targets in the other two *PHT* genes (*PHT3* and *PHT4*), using the specific primers listed in Supplementary Table 2. The amplified fragments were digested using *EcoRI* and *XhoI* restriction enzymes and ligated into the pTRV2 vector also digested with the same enzymes. pTRV1, pTRV2::00, pTRV2::*PHT1/2_*target 1 and pTRV2::*PHT1/2* target 2 were introduced into *Agrobacterium tumefaciens* strain GV3101 pmp90 through electroporation. Transformed strains were cultured in Luria-Bertani (LB) medium supplemented with 10 mM MES and 20 µM acetosyringone at 28°C for 24 h, with appropriate antibiotics. Bacterial cultures were harvested by centrifugation and resuspended in infiltration buffer (10 mM MgCl₂, 10 mM MES, pH 5.7, 100 µM acetosyringone) to an optical density at 600 nm (OD₆₀₀) of 0.5. Suspensions were incubated in darkness at room temperature for 2-3 h prior to infiltration. Equal volumes of *Agrobacterium* cultures containing pTRV1 and pTRV2 derivatives were mixed at a 1:1 ratio before plant inoculation. Tomato seedlings at the two-true-leaf stage were infiltrated with the *Agrobacterium* suspension at the axillary region of the stem using a 1 mL syringe fitted with a needle (Pastor-Fernandez et al., 2024). Fourteen days after inoculation, the plants were used for experiments.

### Plant infestation and assessment of herbivore performance

Five weeks after transplanting tomato plants, the corresponding plants were infested with *S. exigua* larvae. Infestation was achieved by placing five second-instar larvae on the apical leaflet of the third fully expanded leaf (counted from the top). Larvae were confined using a clip cage, and empty clip cages were placed on the leaves of uninfested plants, according to Rivero et al. (2025). After one or two days of herbivory, larvae were removed from the plants and reserved. Damaged apical leaflets were individually harvested and stored at −80°C for subsequent molecular and metabolomic analyses. For undamaged plants, leaf material collection was performed similarly. Following leaf sampling, all larvae corresponding to the same plant were immediately returned to the same plant, to the following leaflet of the same leaf that they were previously feeding on. The entire leaf was enclosed using a mess bag, and larvae were allowed to feed *ad libitum* on the entire leaf. Larvae were removed and their weight and mortality were recorded periodically during the time of the bioassays. Larvae were returned to the same plant, on one leaf above the one they were previously feeding on, to ensure that larvae had enough food during the entire bioassays.

### Bioassay on artificial diet

Caffeoylputrescine (BLDpharm, Germany) and feruloylputrescine (TargetMol, United States) were purchased to test their direct effect on *S. exigua* performance. Both compounds were incorporated, separately or together, into the artificial diet (Hoffman et al., 1966) at a final concentration of 1 µg mL⁻¹ (Rivero et al., 2021). Four different diet formulations were prepared: artificial diet supplemented with caffeoylputrescine (CP), artificial diet supplemented with feruloylputrescine (FP), artificial diet supplemented with both FP and CP (FP+CP), and a control diet supplemented with acidified Milli-Q water (H2Oac, 0.01 % glacial acetic acid), corresponding to the solvent used for compound dissolution. Thirty second-instar *S. exigua* larvae were assigned to each diet treatment and reared under controlled conditions in a growth chamber maintained at 23°C, with a 12-h light and 12-h dark cycle and 50% relative humidity. Larval performance was assessed by monitoring larval weight over a period of 6 days.

### Simulated herbivory treatment

We performed two rows of punctures onto each side of the midvein of the apical leaflet of the third fully expanded leaf (counted from the top) using a fabric pattern wheel. Inmediatley, 5 µL of a 5 mM (Mes-NaOH, pH 6.0) dilution of *S. exigua* oral secretion (OS) was applied to the puncture wounds with a gloved finger. Oral secretions from third instar larvae were collected with a pipette according to (Bricchi et al., 2013), pooled and kept at −20°C until use. The OS quantity was adjusted after several trials.

### RNA extraction and gene expression analysis

Total RNA from fresh leaves was extracted by using the RNeasy Plant Mini Kit (Qiagen) and treated with DNase I (Qiagen) according to the manufacturer’s instructions. First-strand cDNA was synthesized from 0.5 µg using PrimeScript RT reagent kit (Takara). Quantitative PCR was conducted using the SYBR PREMIX EX TAQ (Takara) following the manufacturer’s instructions and a thermal cycler (Bio-RAD, CFX Opus 384) with the following cycling program: 30 sec 95°C, 40 cycles of 5 sec 95°C and 60°C 34 sec, followed by a dissociation process of 95°C 15 sec, 60°C 1 min and 95°C 15 sec. The sequences of the gene-specific primers are shown in Supplementary Table 3. Before normalization, we analyzed the expression of three different tomato housekeeping genes under the different conditions of our study: actin (Solyc03g078400), β-tubulin (Solyc04g081490), and elongation factor 1-α (Solyc06g005060). We used the Normfinder software (https://moma.dk/normfinder-software) to find the optimal normalization gene. According to the results, expression values were normalized using the housekeeping gene elongation factor 1-α (*EF-1α*). The normalized data were further processed by the 2^-ΔΔct^ method (Livak & Schmittgen, 2001).

### Measurement of arginine decarboxylase and ornithine decarboxylase enzymatic activities

Arginine decarboxylase (ADC) and ornithine decarboxylase (ODC) enzymatic activities were determined according to Wu et al. (2012). Briefly, 0.2 g of fresh plant tissue was homogenized in extraction buffer (0.1 M phosphate buffer, pH 5.6, 5 mM EDTA, 1 mM pyridoxal phosphate, 0.01 mM polyvinyl pyrrolidone, 10 mM dithiothreitol (DTT), and 0.43 mM sodium thiosulfate). The homogenate was centrifuged at 12.000 *g* at 4°C for 40 min. A volume of 0.8 mL of the supernatant was mixed with 1 mL of reaction mix (0.1 M Tris-HCl buffer, pH 7.5, 5 mM EDTA, 40 μM pyridoxal phosphate, and 5 mM DTT). To assess enzymatic activity, 0.2 mL of either 25 mM L-Arg (for ADC) or 25 mM L-Orn (for ODC) was added, and the mixture was incubated at 37°C for 60 min. Control reactions in which the substrate was substituted with perchloric acid, were included to determine background activity. Following incubation, 0.5 mL of the sample was mixed with 1 mL of 2 M NaOH and 10 μL benzoyl chloride. The mixture was stirred for 20 s, and incubated for 30 min at 37°C. Subsequently, 2 mL of saturated NaCl and 2 mL of 100% ether were added, followed by a centrifugation at 1500 *g* for 5 min at 4°C. A 1 mL aliquot of the ether phase was collected, evaporated at 50°C in a water bath, and the remainder was dissolved in 1 mL of 100% methanol. The benzoylated polyamine derivatives were quantified at 254 nm using a GeneQuant 1300 spectrophotometer (Harvard Bioscience Inc., Holliston, MA, USA). Enzyme activity was calculated based on the increase in absorbance relative to control samples. One unit of enzymatic activity was defined as an increase of 1.0 absorbance unit at 254 nm during 1 min.

### Determination of free polyamines content

The content of free putrescine (Put), spermidine (Spd) and spermine (Spm) in leaf extracts was measured as described in Ariz et al. (2013) with some modifications. Fresh plant tissue was ground with liquid nitrogen and homogenized in 5% aqueous HClO_4_ (w: w) at a ratio of 10:1 (v : w), supplemented with an internal standard solution (2 mM 1,6-hexanediamine) at 0.1:1 (v : w). Homogenates were shaken, incubated for 1 h at 4°C, and centrifuged at 15 000 *g* for 15 min at 4°C. Aliquots (0.2 mL) of the supernatant were mixed with 0.4 mL of 3 M aqueous Na_2_CO_3_ and 0.4 mL of 0.12 M dansyl chloride in acetone, followed by incubation at 60°C in darkness for 1 hour. To stop the reaction, 0.1 mL of 0.87 M proline were added and the mixture was further incubated for 30 min. Dansylated polyamines were extracted using 5 mL of ethyl acetate, centrifuged at 3000 *g* for 5 min at room temperature and the organic phase was collected and evaporated under reduced pressure at 40°C. The residue was resuspended in 0.5 mL of methanol and filtered through a 0.45 μm nylon filter. Free polyamine separation and quantification were carried out by high-performance liquid chromatography (HPLC) using a Waters system (616 pump, 717 Plus Autosampler, and 2475 Fluorescence Detector; Waters, Milford, MA, USA). A Tracer Excel 120-ODS-A column (3 μm, 4.6 × 150 mm; Teknokroma, Barcelona, Spain) maintained at 30°C was used as the stationary phase for separation. The mobile phase consisted of solvent A (water) and solvent B (methanol), with a constant flow rate of 0.5 mL min⁻¹. An increasing concentration gradient was applied, solvent B going from 58% to 100% over 44 min, held for 4 min, then returned to 58% over 3 min, and equilibrated for an additional 3 min. Fluorescence detection was performed at an λ_ex_ of 350 nm and an λ_em_ of 515 nm. Quantification of polyamines was performed using an internal standard and calibration curves prepared with authentic standards of Put, Spd, and Spm.

### Determination of hydrogen peroxide content

Hydrogen peroxide was determined from 200 mg of leaf homogenized in cold with 2 mL of 0.1% TCA. The samples were centrifuged at 4°C for 15 min at 17100 *g*. The H_2_O_2_ concentration was determined in the supernatant using a modification of the ferric thiocyanate method (Sagisaka et al., 1976). Briefly, 0.2 g of fresh tomato leaf tissue was homogenized in 2 mL of a 0.1% trichloroacetic acid. Three technical replicates were performed for each biological replicate, and samples were analyzed using a ClarioStar spectrophotometer. A hydrogen peroxide solution was used to setup a standard curve.

### Determination of γ-aminobutyric acid (GABA) content

Approximately 5 mg of sample were weighed and dissolved in 1.75 mL of 1 M borate buffer (pH 9) and 0.75 mL of methanol. Subsequently, 30 µL of diethyl ethoxymethylenemalonate were added, and the reaction mixture was incubated at 70°C for 2 h. After incubation, the solution was diluted 10-fold in a mixture of methanol and 10% formic acid. The resulting solution was directly injected into the liquid chromatograph together with calibration standards. Quantification was performed based on the exact amount of sample weighed. Liquid chromatography-mass spectrometry (LC-MS) analysis was carried out by the Nucleus Platform (University of Salamanca) using a Thermo Vanquish system coupled to a Thermo Orbitrap Q Exactive Focus mass spectrometer. Chromatographic separation was achieved using a Phenomenex Luna Omega C18 column (50 mm × 2.1 mm, with internal diameter 1.6 µm particle size). The mobile phases consisted of water with 0.1% formic acid (solvent A) and methanol (solvent B), with a flow rate of 0.4 mL min⁻¹. The gradient program was as follows: 30% B at 0 min, increased to 60% B at 11 min, to 100% B at 12 min, held at 100% B until 15.5 min, returned to 30% B at 16 min, and equilibrated at 30% B until 20 min.

### Untargeted metabolomics by liquid chromatography and electro-spray ionization (LC-ESI) mass spectrometry

Metabolites were extracted from 20 mg of freeze-dried leaves using MeOH:H2O (10:90) containing 0.01% of HCOOH, as described by Rivero et al. (2021). Extracts were injected into an Acquity ultra-performance liquid chromatography system (UPLC; Waters, Mildford, MA, USA), interfaced with a hybrid quadrupole time-of-flight equipment (Q-TOF-MS Premier, Waters, Mildford, MA, USA). Analytes were eluted with an aqueous methanol gradient containing 0.01% HCOOH. Solvent gradients and further chromatographic conditions were performed as previously described (Agut et al., 2014). LC separation was performed using an ultra-performance liquid chromatography (UPLC) Kinetex C18 analytical column with a 5 μm particle size, 2.1 × 100 mm (Phenomenex, Madrid, Spain). To accurately identify the signals detected, a second fragmentation function was introduced into the TOF analyser. This function was programmed in a t-wave ranging from 5 to 45 eV to obtain a fragmentation spectrum of each analyte (Agut et al., 2014; Gamir et al., 2014). Ornithine and arginine were identified by using commercial standards. To specifically identify phenolamides, we focused our analysis in hydroxycinnamic acid conjugates of polyamines, using the retention times and mass fragmentation patterns of reference compounds for caffeoylputrescine and feruloylputrescine, as well as the characteristic mass fragmentation of feruloyldicoumaroylspermidine.

### Statistical analysis

Statistical analyses were performed using R software (R version 4.4.3) in RStudio (v1.4.1564). Data normality and homogeneity of variances were assessed using the Shapiro-Wilk and Levene’s tests, respectively. Kaplan-Meier survival curves were generated for herbivore performance assays using the *survival* and *survminer* packages. One-way ANOVA was used to analyze larval weight datasets, with pairwise comparisons conducted using Student’s *t*-test (*P* ≤ 0.05). Two-way ANOVA models were applied to evaluate the effects of Trichoderma treatment and genotype on larval weight data; and to evaluate the effects of herbivory (or herbivory mimic) and Trichoderma treatment on gene expression and metabolite. Tukey’s test (*P* ≤ 0.05) was used for overall comparisons among treatment groups. All graphical outputs were generated using the *ggplot2* package. All experiments were repeated at least twice, with similar results.

### Accession numbers

Sequence data used in this article can be found in the Solgenomics platform (https://solgenomics.net/) under the following ITAG4 accession nos.: Solyc10g054440; Solyc11g071480; Solyc04g082030; Solyc03g098300; Solyc12g038970; Solyc11g068540; Solyc01g091160; Solyc01g091170; Solyc05g005710; Solyc03g007240.3; Solyc05g010420; Solyc8g075710; Solyc05g013440; Solyc09g075940; Solyc08g079430.2; Solyc01g087590; Solyc01g087590; Solyc11g071470; Solyc11g071480; Solyc06g074710; Solyc07g008390; and Solyc06g005060.

## AUTHOR CONTRIBUTIONS

IF and SCT contributed equally. IF and SCT performed most of the experiments and analyses, and contributed to design research, data interpretation and manuscript writing. BRR designed and carried out VIGS bioassays and data interpretation. PLG assisted with greenhouse bioassays, and RT-qPCR and enzymatic activity analyses. AIGH assisted with bioassays with the RNAi line. AC performed quantification of free polyamines and data interpretation. VF performed LC-ESI mass spectrometry analyses and contributed to data interpretation. MJP contributed to design the research, data interpretation and manuscript writing. AMM designed and supervised research and wrote the manuscript.

## SUPPLEMENTARY DATA

**Supplementary Figure 1**: Impact of *Trichoderma harzianum* root colonization on tomato growth.

**Supplementary Figure 2**: Schematic illustration of polyamine metabolism in plants.

**Supplementary Figure 3**: Impact of *Trichoderma harzianum* root colonization and *Spodoptera exigua* herbivory on the expression of *ARGINASE 1*.

**Supplementary Figure 4**: Effect of *Trichoderma harzianum* root colonization and *Spodoptera exigua* herbivory on *PHT3* and *SHT* expression; and identification and quantification of putrescine- and spermidine-derived phenolamides.

**Supplementary Table 1**: Results (*F* and *P* values) from 1-Way and 2-Way ANOVAs for the different variables presented in Figures 2-5 in main text.

**Supplementary Table 2**: *SlPHT2* gene target regions and primer sequences for Virus Induced Gene Silencing (VIGS)

**Supplementary Table 3**: Specific primers used for RT qPCR

## FUNDING

This research was supported by the European Union (ERC, ERC-2023-COG: 101124883 MIMIR), by the grants PID2021-128318OA-I00 and PID2024-162058OB, funded by MICIU/AEI/10.13039/ 501100011033 and FEDER, UE.

## Notes

### Competing Interest Statement

The authors have declared no competing interest.

### Summary of Updates

Minor changes have been made, including the grouping of figures and the addition of new analyses. Some typographical errors have also been corrected. The main conclusions of the paper remain unchanged.

